# Hippocampal-neocortical interactions sharpen over time for predictive actions

**DOI:** 10.1101/483115

**Authors:** Nicholas C. Hindy, Emily W. Avery, Nicholas B. Turk-Browne

**Affiliations:** Psychological and Brain Sciences, University of Louisville, Louisville, KY 08544; Psychology, Yale University, New Haven, CT 08544

## Abstract

When an action is familiar, we are able to anticipate how it will change the state of the world. These expectations can result from retrieval of action-outcome associations in the hippocampus and the reinstatement of anticipated outcomes in visual cortex. How does this role for the hippocampus in action-based prediction change over time? We used high-resolution fMRI and a dual-training behavioral paradigm to examine how the hippocampus interacts with visual cortex during predictive and nonpredictive actions learned either three days earlier or immediately before the scan. Just-learned associations led to comparable background connectivity between the hippocampus and V1/V2, regardless of whether actions predicted outcomes. However, three-day-old associations led to stronger background connectivity and greater differentiation between neural patterns for predictive vs. nonpredictive actions. Hippocampal prediction may initially reflect indiscriminate binding of co-occurring of events, with action information pruning weaker associations and leading to more selective and accurate predictions over time.

## Introduction

As you open the door to a familiar room, you are able to anticipate that specific objects that will come into view. A neural source of such predictions may be pattern completion in the hippocampus^1–3^. Repeated experience and interaction allows associative learning mechanisms in the hippocampus to bind recurring patterns of objects and actions over space and time^4,5^. Once these links are formed, making an action in response to a familiar cue may prompt the hippocampus to retrieve a conjunctive representation of past events. These representations could contain information about the cue and action, but additionally the yet-to-occur sensory consequences of the action. These retrieved consequences could in turn get reinstated via feedback to sensory systems — a form of memory-based predictive coding of action outcomes.

A recent study provided suggestive evidence for this mechanism, discovering a link between pattern completion in the hippocampus and predictive coding in visual cortex^3^. Participants were trained behaviorally on cue-action-outcome sequences: in response to a visual cue, they chose between two manual actions that were either predictive or nonpredictive of the visual outcome that next appeared. The day after training, participants were scanned with fMRI while performing the same task with the pre-learned associations. However, on a subset of trials in the scanner, the outcome stimulus was omitted and replaced by a blank screen. For predictive actions (i.e., actions that determine an outcome given a cue), multivariate pattern decoding revealed that the hippocampus represents the full cue-action-outcome sequence and that this is related within and across participants to evidence of the same outcome in early visual cortex (EVC), as measured with a separate classifier trained on outcome-only trials. No reliable decoding effects were obtained in either the hippocampus or EVC for nonpredictive actions (i.e., actions that did not determine which outcome appeared after a cue).

This role for the hippocampus in action-based predictive coding can be interpreted in two ways. First, it could be explained in terms of the unique computational repertoire of the hippocampus, with processes like multimodal binding and pattern completion serving an important function in prediction regardless of the more canonical role of the hippocampus as a memory system^6,7^. Second, the hippocampus may have been involved because the associations had been learned very recently, and this role would diminish over time with the neocortex playing a more autonomous role as a result of systems consolidation^8–10^ (cf. ^11^). The previous study was unable to disentangle these possibilities because all associations had been learned at a fixed time in the past. Therefore, in the current study we tested the role of the hippocampus in action-based prediction over two timescales. Moreover, whereas the previous study was suggestive of hippocampal-neocortical interactions during prediction, this was based on the correlation of static information present in both systems. Here we more directly measured these interactions using a “background connectivity” approach that quantifies the temporal dynamics and covariance of the hippocampus and EVC^12,13^. We hypothesized that background connectivity between the hippocampus and EVC would depend on both the lag between training and scanning and the predictiveness of actions, and that this would relate to the representational contents of these areas.

Participants learned cue-action-outcome sequences in a first training session three days before an fMRI scan and in a second training session immediately before the scan (Fig. 1A). Separate sets of cues and outcomes were used in each training session and actions were either predictive or nonpredictive of outcomes depending on the cue (Fig. 1B). For predictive actions, one outcome reliably followed the cue after a left button press and a different outcome reliably appeared after a right button press; explicit memory of predictable outcomes was at ceiling on verbal tests administered during each training session and before and after the fMRI scan (Fig. 1C). For nonpredictive actions, the two outcomes followed the cue with equal probability when either the left or right button was pressed. After both training sessions, participants performed the same task in the fMRI scanner, with stimuli from the two training sessions presented separately in alternating runs, and cues with predictive vs. nonpredictive actions presented separately in alternating blocks within each run type. Background connectivity was calculated for each of these blocks and then collapsed within condition, resulting in four key measures of hippocampal-EVC interaction: 3-day vs. no delay learning of predictive vs. nonpredictive actions.

**Figure 1.**
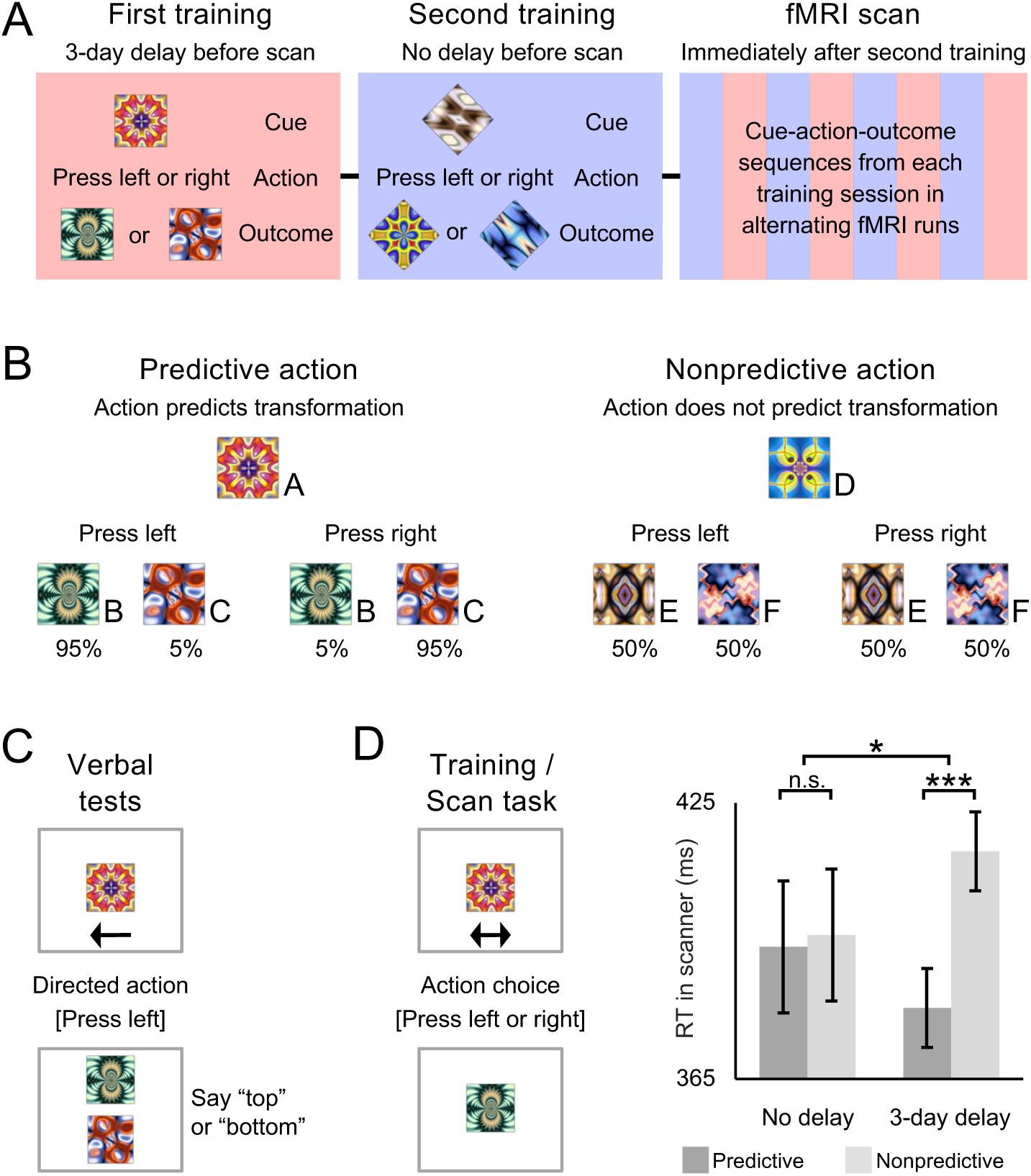
Design and behavior. **(A)** The first training session was proctored three days before the fMRI scan, while the second training session was proctored immediately before the scan. To reduce interference, fractal stimuli in each session were masked to be either squares or diamonds in shape. The fMRI scan included 4 runs for each of the two training sets. **(B)** For predictive actions, given the cue, one outcome appeared with 95% probability when the left button was pressed and the other with 95% probability when the right button was pressed. For nonpredictive actions, given the cue, either outcome appeared with 50% probability when the left or right button was pressed. **(C)** In verbal tests outside the scanner, participants spoke aloud either “top” or “bottom” to indicate which outcome was expected given the cue and action. **(D)** For each trial in the fMRI scanner, participants chose to press either a left-hand or right-hand button to replace a cue with an outcome. While choice RT was similar for predictive and nonpredictive actions immediately after training, it was faster for predictive vs. nonpredictive actions after a 3-day delay. Error bars indicate ±1 SEM of the difference between predictive and nonpredictive actions at each timescale. \*\*\**p* <.001; \**p* <.05.

## Results

### Verbal tests of explicit memory

To verify that predictive actions had been learned during training and remembered across the delay, participants were required to be 100% accurate in identifying expected outcomes of predictive actions in verbal outcome-identification tests outside of the fMRI scanner. Participants who did not reach this accuracy criterion on each test were excluded from the fMRI scan. Thus, by definition, all 24 scanned participants reached perfect accuracy. Two additional participants completed training but did not participate in the fMRI scan because of accuracy less than 100% even after repeating a pre-scan test.

### Choice RT for predictive and nonpredictive actions

Throughout training and in the scanner, we measured choice response time (RT) as the time it took for participants to press the left or right button in response to a cue. During training sessions outside of the scanner, choice RT did not differ among the conditions (all *p*s >.26). The lack of a timescale difference between training sessions is not surprising, as these conditions were equivalent at this point in the study. However, in the scanner, we observed a reliable interaction between timescale and predictiveness (*F*(1, 22) = 5.49, *p =.*03; Fig. 1D). For no-delay sequences, choice RT was comparable for predictive and nonpredictive actions (*t*(23) = 0.18, *p =.*86), whereas for 3-day delay sequences, choice RT was faster for predictive vs. nonpredictive actions (*t*(23) = 3.96, *p* <.001).

### Stimulus-evoked responses

A general linear model (GLM) containing finite impulse response (FIR) basis functions was used to estimate evoked blood-oxygen level-dependent (BOLD) activity in the hippocampus and EVC (Fig. 2A). Stimulus-evoked activity for each condition was averaged within block to capture the peak response. Although activity in the hippocampus was marginally reduced for both predictive and nonpredictive actions after the 3-day delay (*F*(1, 22) = 4.09, *p* =.05), no other main effects or interactions were observed in either ROI (*p*s >.35).

**Figure 2.**
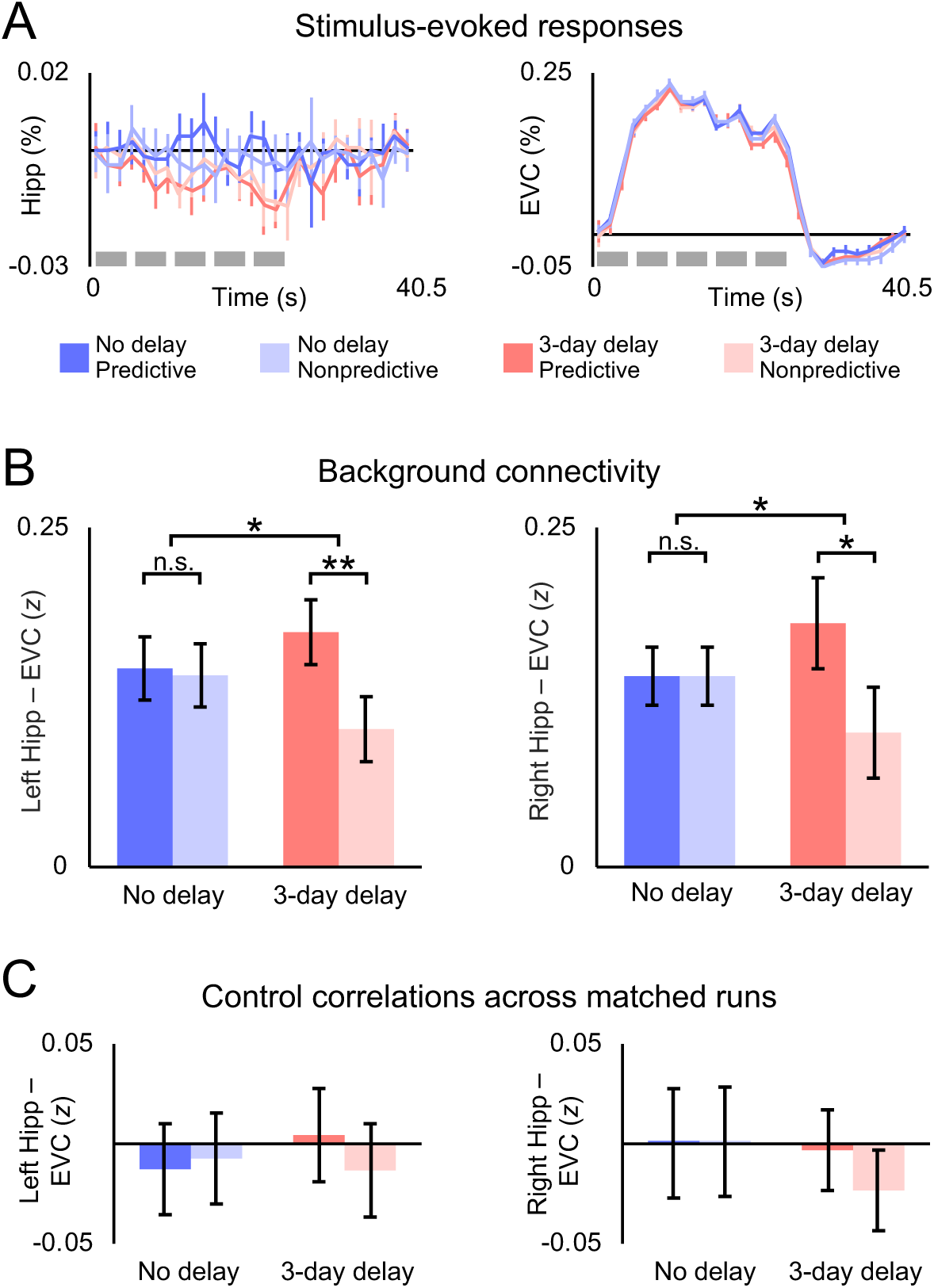
Stimulus-evoked responses and background connectivity. **(A)** Within each ROI, stimulus-evoked BOLD activity was similar during blocks of predictive and nonpredictive actions, both immediately after training and three days after training. Each block contained five trials (gray bars). **(B)** For both left and right the hippocampus, background connectivity with EVC was stronger for predictive vs. nonpredictive actions three days after training but not immediately after training. **(C)** Control correlations across runs with matching stimuli and timing did not differ between any of the conditions. Error bars indicate ±1 SEM of the difference between predictive and nonpredictive actions at each timescale. \*\**p* <.01; \**p* <.05.

### ROI background connectivity

Task-specific background connectivity between the hippocampus and EVC was measured after removing stimulus-evoked activity and confounding variables through linear regression in a multistep procedure published previously^12,14–17^. We first used a GLM to regress out white matter and ventricle activity along with motion parameters from preprocessing, and then used FIR basis functions to capture and remove the average timing and shape of the hemodynamic response in each voxel in a data-driven way. Background connectivity was measured as correlations in the residual timeseries of each ROI. There were no differences across hemispheres in background connectivity between the hippocampus and EVC (*p*s >.61). Critically, we observed a reliable interaction between timescale and predictiveness (*F*(1, 23) = 8.28, *p =.*008; Fig. 2B). This interaction was driven by a reliable difference between predictive and nonpredictive actions for sequences learned three days before the fMRI scan (*t*(23) = 2.90, *p =.*008), with no hint of an effect of predictiveness on sequences learned immediately before the scan (*t*(23) = 0.12, *p* =.90). When predictive and nonpredictive events were separately compared across delay conditions, enhanced background connectivity over time for predictive actions was not significant (*t*(23) = 1.67, *p* =.11), while diminished background connectivity over time for nonpredictive actions was significant (*t*(23) = 2.34, *p* =.03).

### Control correlations across matched runs

The goal of background connectivity is to remove stimulus-evoked responses to isolate idiosyncratic fluctuations that reveal how experimental conditions modulate functional connectivity. To verify that the residualizing approach above was effective, we performed an across-run control analysis^12^. Each training condition was tested in two runs that used the same block order and the same cue-stimulus-outcome sequences. If the key findings above were confounded by unmodeled stimulus-evoked responses, the residual activity in the hippocampus in one run should be correlated with the residual activity in EVC in the other run. However, there were no reliable interactions or main effects when computing connectivity across runs (*p*s >.17; Fig. 2C); moreover, connectivity was reliably lower for each condition when it was calculated across vs. within run (*p*s <.01).

### Voxelwise background connectivity

To what extent are timescale and predictiveness differences in background connectivity between the hippocampus and EVC specific to these regions vs. widespread in the brain? To assess the anatomical specificity of the effects, we performed exploratory analyses using the residual timeseries from bilateral hippocampus and EVC ROIs (Fig. 3A) to calculate background connectivity with all voxels in the partial volume collected for each participant. After registering these correlation maps to MNI space, we conducted nonparametric randomization tests of their reliability across participants. Immediately after training, predictiveness did not reliably modulate background connectivity anywhere in the partial volume when the hippocampus or EVC served as the seed (Fig. 3B). Conversely, for sequences learned three days before the scan, several clusters showed reliably greater background connectivity with each seed during blocks of predictive vs. nonpredictive actions (Fig. 3C). Specifically, predictiveness enhanced the background connectivity of the hippocampus with left (−9, −87, −13) and right (19, −91, −10) occipitotemporal cortex and left (−20, 6, −3) and right (25, 18, −4) putamen, and enhanced the background connectivity of EVC with anterior (−30, −12, −27) and posterior (−21, −42, −7) left hippocampus (bilateral at uncorrected threshold), left parahippocampal gyrus (−17, −52, 2), and left posterior cingulate cortex (−9, −55, 14). At each timescale, no voxels showed stronger background connectivity with the hippocampus or EVC for nonpredictive actions.

**Figure 3.**
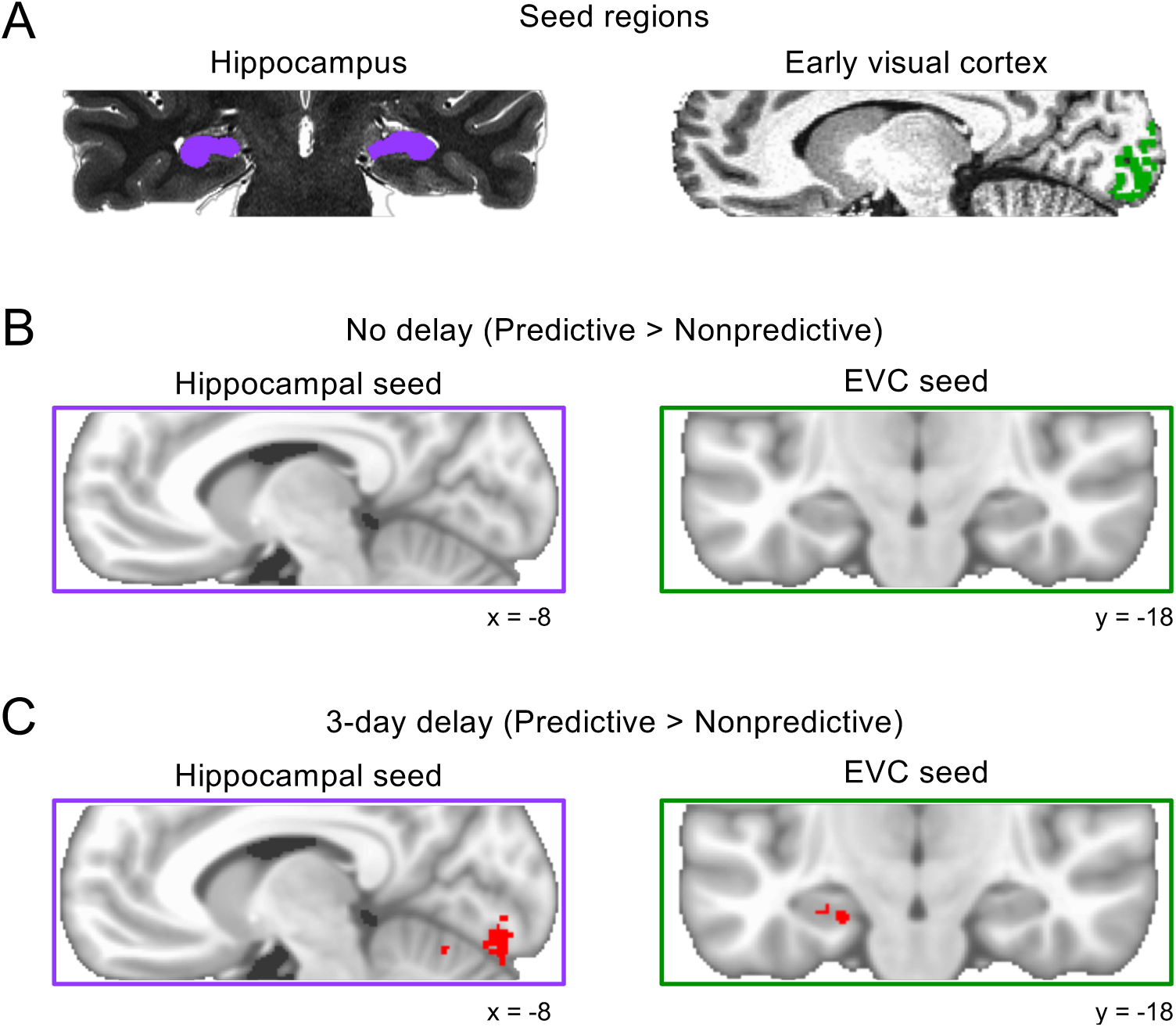
Voxelwise background connectivity. **(A)** Hippocampal and EVC seed regions were defined on high-resolution anatomical scans. **(B)** Predictiveness did not reliably modulate hippocampal or EVC background connectivity anywhere in the field of view immediately after training. **(C)** However, we observed several clusters of enhanced background connectivity after the 3-day delay, including bilateral V1 and V2 for the hippocampal seed, and anterior and posterior left hippocampus for the EVC seed. Contrasts are visualized on the MNI152 template and corrected at *p* <.05 (TFCE) for the partial volume of functional scans outlined by purple/green boxes.

**Figure 4.**
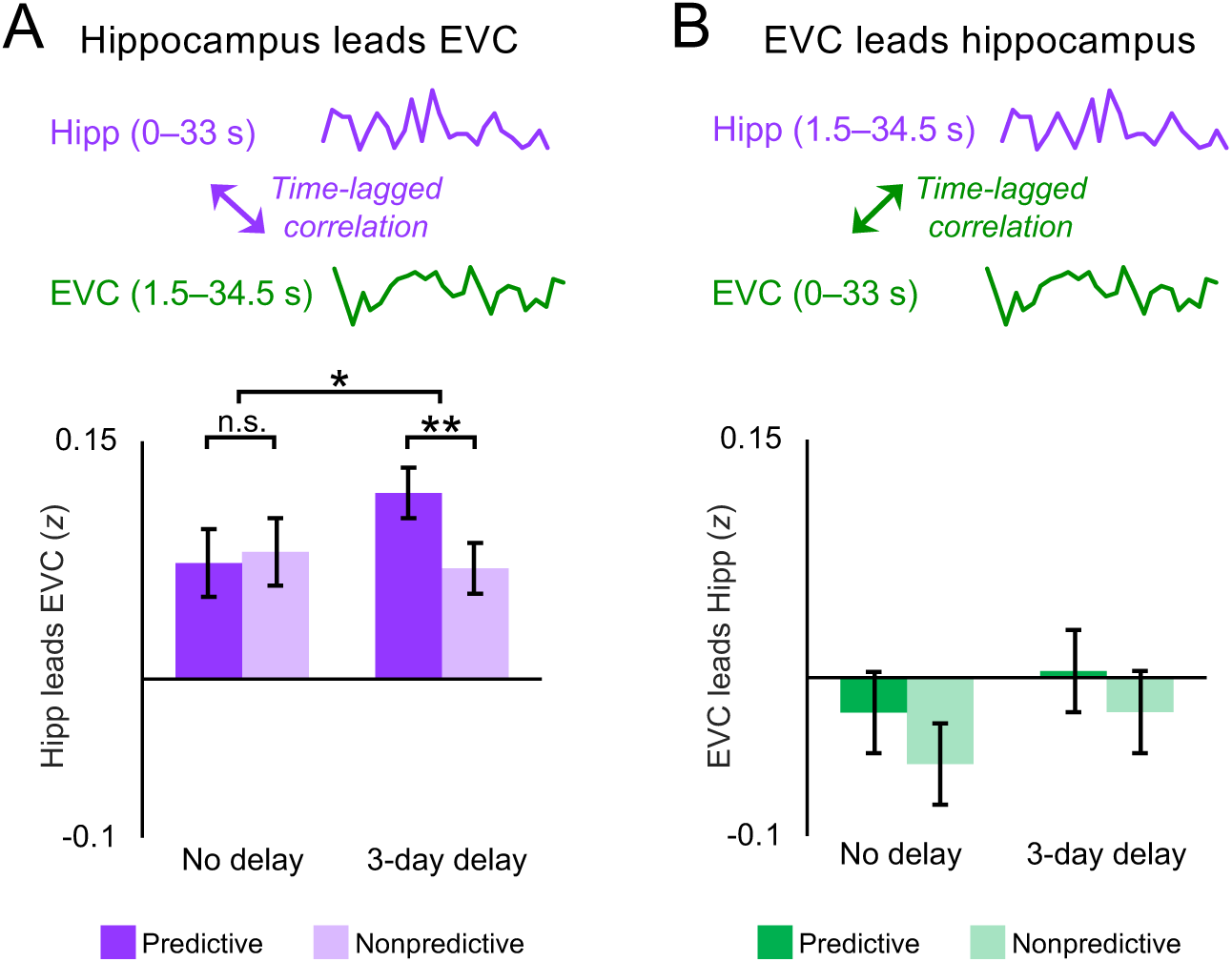
Time-lagged background connectivity. **(A)** Hippocampus leads EVC: For each block we computed the correlation between earlier background activity in the hippocampus (0–33 s) with later background activity in EVC (1.5–34.5 s); a similar pattern of results emerged as for non-shifted background connectivity. **(B)** EVC leads hippocampus: For each block we correlated earlier background activity in EVC (0–33 s) with later background activity in the hippocampus (1.5–34.5 s); there were no reliable differences among conditions. Error bars indicate ±1 SEM of the difference between predictive and nonpredictive actions at each timescale. \*\**p* <.01; \**p* <.05.

### Time-lagged background connectivity

Background connectivity between the hippocampus and visual cortex during predictive action is agnostic to the direction of the interaction. Such questions can only be addressed definitively with techniques that allow for causal interventions. Moreover, the slow sampling rate of fMRI and the temporal autocorrelation of BOLD activity severely limit the analysis of temporal dynamics. Nevertheless, it is possible to test whether there exists any evidence for a temporal asymmetry in the signals between these regions^3^ that would be consistent with processing in one region preceding the other. Specifically, we hypothesized that insofar as the hippocampus is relying on learned predictiveness to reinstate expected outcomes in visual cortex, the activity in the hippocampus at one time point should predict activity in visual cortex at the next time point, at least more than the reverse. Indeed, we were able to replicate the main timescale by predictiveness interaction reported above when EVC was lagged by one time point with respect to the hippocampus (*F*(1, 22) = 4.77, *p =.*04). This interaction reflected a reliable difference in background connectivity between predictive and nonpredictive actions for sequences learned three days before the fMRI scan (*t*(23) = 2.98, *p =.*007) and not for sequences learned immediately before the scan (*t*(23) = −0.33, *p =.*74). Critically, using nonpredictive blocks as the baseline controls for the possibility that BOLD activity merely peaks later in visual cortex than the hippocampus. In contrast, no such interaction was found when the hippocampus was lagged with respect to EVC (*t*(23) = 0.04, *p =.*84), with no differences between predictive and nonpredictive actions at either timescale (*p*s >.21).

### Multivariate pattern similarity

We have shown that background connectivity between the hippocampus and EVC strengthens over time for predictive relative to nonpredictive actions. How does this relate to the information represented in each ROI? Specifically, we hypothesized that greater connectivity for predictive actions after three days should be accompanied by greater information about expected outcomes. We tested this by measuring the neural similarity between sequences in which the same cue appeared but was followed by different outcomes (Fig. 5A). Insofar as these overlapping sequences are more differentiated after three days vs. immediately after training, it would imply that the actions led to a stronger and/or clearer prediction of the outcome. To calculate pattern similarity, we correlated spatial patterns of parameter estimates in the hippocampus and EVC obtained from an event-related GLM, as a function of which cue was presented, whether it was associated with predictive vs. nonpredictive actions, and at what timescale it was learned.

**Figure 5.**
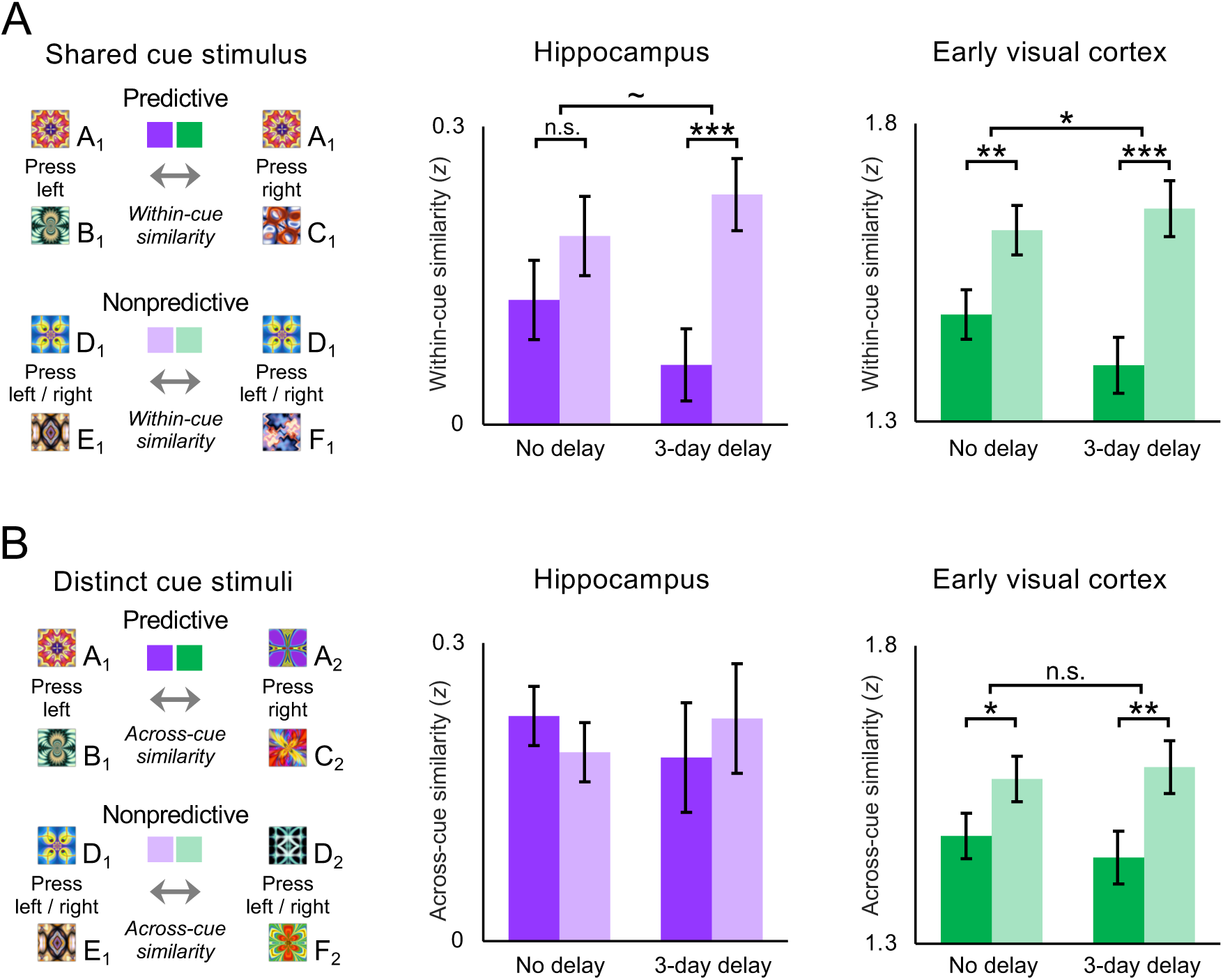
Multivariate pattern similarity. **(A)** Within-cue pattern similarity was measured as the correlation across voxels for sequences that shared the same cue but contained different outcomes. Hippocampal and EVC within-cue similarity was lower for predictive than nonpredictive actions after a 3-day delay, compared to no delay. **(B)** Across-cue pattern similarity was measured between sequences with non-overlapping stimuli. Delay interval did not modulate across-cue similarity in either the hippocampus or EVC. Error bars indicate ±1 SEM of the difference between correlations for predictive and nonpredictive actions at each timescale. \*\*\**p* <.001; \*\**p*<.01; \**p*<.05; ∼*p* =.05.

Consistent with a prior study^3^, neural representations of the two sequences associated with each predictive cue were less similar to one another than those of each nonpredictive cue in both the hippocampus (*F*(1, 23) = 17.77, *p* <.001) and EVC (*F*(1, 23) = 32.16, *p* <.001), suggesting that predictive actions helped disambiguate action outcomes in these regions. Importantly, this differentiation effect was modulated by delay condition in both the hippocampus (*F*(1, 23) = 4.26, *p* =.05) and EVC (*F*(1, 23) = 5.36, *p* =.03). In the hippocampus, pattern similarity was reliably reduced for predictive vs. nonpredictive cues and actions trained three days before the scan (*t*(23) = 4.73, *p* <.001), but did not differ for actions trained immediately before the scan (*t*(23) = 1.61, *p =.*12). In EVC, despite a reliable interaction, the difference in pattern similarity for predictive and nonpredictive actions was reliable both after the 3-day delay (*t*(23) = 5.60, *p* <.001) and immediately after training (*t*(23) = 3.42, *p =.*002). Unlike background connectivity, pattern similarity did not significantly differ between immediate and 3-day delay conditions within just predictive or just nonpredictive actions in either ROI (*ps* >.10).

Do predictive events become more neurally distinct than nonpredictive events specifically when they share a cue (and thus initially overlap), or do they become more neurally distinct in general? To test whether differences in pattern similarity extend to non-overlapping events, we measured similarity between sequences with different cues (Fig. 5B). In the hippocampus, there was no difference in pattern similarity between predictive vs. nonpredictive actions when both the cues and the outcomes were distinct (*F*(1, 23) = 0.00, *p =.*97). Conversely, this difference was reliable in EVC (*F*(1, 23) = 17.24, *p* <.001), with reduced similarity between sequences with predictive vs. nonpredictive actions. However, unlike the differentiation effect between overlapping sequences, across-cue similarity did not interact with delay condition in either the hippocampus (*F*(1, 23) = 1.74, *p =.*20) or EVC (*F*(1, 23) = 0.96, *p =.*34). Across-cue pattern similarity also did not reliably differ between immediate and 3-day delay conditions within just predictive or just nonpredictive actions in either ROI (*p*s >.16).

## Discussion

Using high-resolution fMRI and a multi-session training paradigm, we examined how functional interactions between the hippocampus and early visual cortex change over the early periods of a memory. Results build upon recent evidence of a link between hippocampal pattern completion and predictive coding in visual cortex^3^, but suggest that the role of the hippocampus in visual prediction depends on the age of the knowledge on which the prediction was based. Specifically, interactions between the hippocampus and visual cortex became weaker for nonpredictive actions (and relatively stronger for predictive actions) three days after learning compared to immediately after learning. Over the same timescale, predictive actions led neural representations in these regions to become more differentiated for sequences with overlapping stimuli. Hippocampal prediction may be based at first on indiscriminate binding of co-occurring stimuli, with time and offline processing leading to gradual pruning of weaker associations, in this case associations without informative actions.

Immediately after training, hippocampal-neocortical interactions were the same for predictive and nonpredictive actions. At first glance, the absence of a difference in background connectivity between these conditions may appear to be odds with previous MVPA findings in which classifier accuracy was at chance in both the hippocampus and EVC for nonpredictive actions while above chance for predictive actions^3^. Critically, however, background connectivity and multivariate pattern decoding are differentially sensitive to prediction in this task. Specifically, although participants cannot accurately predict outcomes of nonpredictive actions, they may nonetheless *inaccurately* predict outcomes. For example, the less predictable transitions for these cues may encourage hypothesis testing or other attempts to continue learning, or they may be predicting both outcomes associated with the cue (which each still co-occur 50% of the time, far higher than any other outcome). Less differentiated patterns in visual cortex may in fact reflect less differentiated neural predictions, as opposed to a lack of prediction. Likewise, in any of these cases, a multivariate classifier will seek evidence of the correct outcome, and so performance will be at chance on average. However, to the extent that background connectivity between the hippocampus and visual cortex reflects the process of prediction, whether accurate or inaccurate, it may be enhanced for both predictive and nonpredictive actions.

Voxelwise background connectivity, corrected for multiple comparisons across the acquired field of view, converged with the ROI findings. Immediately after training, predictive actions did not reliably modulate voxelwise background connectivity with seeds for either the hippocampus or EVC. However, after a 3-day delay, predictive actions significantly modulated hippocampal background connectivity with voxel clusters in V1 and V2, as well as EVC background connectivity with voxel clusters in the hippocampus. While voxelwise background connectivity largely overlapped with the *a priori* ROIs, a few other interesting findings emerged including enhanced hippocampal background connectivity for predictive actions with the left and right putamen and with object-selective visual areas in posterior fusiform and lateral occipital cortex. The putamen is especially intriguing because it has frequently been linked with action selection^18^ and with offline motor-sequence learning^19^. Moreover, this finding converges with previous MVPA findings for action decoding in the right putamen^3^.

Along with background connectivity, multivariate pattern similarity within the hippocampus and visual cortex depended on the combination of predictiveness and delay interval. In the hippocampus, consistent with hippocampal models of episodic memory that emphasize the importance of representational overlap for neural differentiation^20,21^, reduced pattern similarity for predictive relative to nonpredictive actions was observed only between sequences that shared the same cue stimulus. In early visual cortex, predictive actions led to more distinctive neural patterns at each time scale for both overlapping sequences (that shared a cue stimulus) and non-overlapping sequences (in which both the cue and outcome differed). However, for non-overlapping sequences, reduced EVC pattern similarity for predictive vs. nonpredictive actions was the same after a 3-day delay as immediately after training. In contrast, delay condition significantly modulated the effect of predictiveness on EVC pattern similarity between overlapping sequences. Thus, the passage time modulated neural differentiation effects in EVC in the same way as predictably observed in the hippocampus, further linking these regions together.

Explicit memory, measured through verbal outcome-identification tests, did not differ between predictive and nonpredictive actions either during the training sessions or immediately before or after the fMRI scan. Yet, three days after training, participants were significantly faster in making predictive actions than nonpredictive actions in the scanner. Dissociations in background connectivity and multivariate pattern similarity may thus reflect changes across time in the perceptual fluency of cue-action-outcome sequences, as opposed to changes in the explicit memory of stimulus outcomes. Critically, however, perceptual fluency in the action-based prediction of upcoming stimuli may be distinct from perceptual fluency in the recognition of static visual stimuli^22,23^. While skill acquisition in purely perceptual tasks such as texture discrimination^24^ and visual contour integration^25^ may be independent of the hippocampus, hippocampal function is necessary for learning arbitrary associations among stimuli^26^. Notably, however, the statistical learning required for action-based prediction may involve different pathways within the hippocampus than other forms of hippocampally dependent learning^5,26^.

Interactions between timescale and predictiveness update consolidation-based models of memory and offline learning that posit a reduced role for the hippocampus over time^8,27^, suggesting that reduced hippocampal involvement may apply first to nonpredictive associations. That is, predictive action may provide a mechanism for prioritizing which representations are either strengthened through synaptic potentiation or weakened through synaptic depression during periods of offline rest^28,29^. Activity-dependent synaptic potentiation and depression may in turn be mediated by offline replay within the hippocampus^30,31^ and between the hippocampus and neocortex^32,33^. By transforming noisy recent associations into sparser remote associations, this offline processing may increase the efficiency and utility of hippocampal associations over time^34,35^. Ultimately, sparser hippocampal representations may increase the signal-to-noise of the hippocampal-neocortical interactions during action-based prediction.

By revealing consolidation-related effects on visual prediction, our findings further develop the link between hippocampal representation^2,36^ and models of predictive coding in visual cortex^37,38^. While feedback across layers of visual cortex may be sufficient to fill-in adjacent elements of a sequence or scene, top-down connections such as from the hippocampus may be needed to simultaneously predict multiple elements in a sequence^39,40^ and to make predictions based on prior co-occurrence and arbitrary associations^3,7^. Indeed, time-lagged background connectivity here converges with previous MVPA findings in which sequence information in the hippocampus temporally preceded outcome information in EVC during mnemonic prediction^3^. When calculating temporal asymmetries in the interaction of hippocampus and EVC, we accounted for potentially confounding differences in the shape and timing of the hemodynamic response by removing evoked activity with data-driven FIR models and by conducting control analyses with nonpredictive blocks and across-run correlations. The timescale by predictiveness interaction observed for background connectivity was preserved when hippocampal background activity was shifted earlier to lead EVC, while it was eliminated when hippocampal background activity was shifted later to trail EVC. Although the causal direction of the relationship between the hippocampus and EVC cannot be established with correlational measures such as fMRI, converging data across this experiment and a previous study^3^ are at least consistent the hippocampus reinstating expected outcomes in visual cortex.

In sum, interactions between the hippocampus and early visual cortex, and representations in these areas, strengthen over time for predictive actions relative to nonpredictive actions. Hippocampal prediction may occur by default, based at first on indiscriminate binding of co-occurring of stimuli. Time and offline processing may gradually prune weaker associations, in this case ones without informative actions, so that hippocampal reinstatement becomes increasingly specific to predictive events.

## Methods

### Participants

Twenty-four individuals (19 female, aged 18–33 years) from the Princeton University community participated in the study. Each participant was right-handed and had normal or corrected-to-normal vision. Two additional participants completed the training sessions but did not participate in the fMRI component of the experiment due to below-criterion accuracy on verbal outcome-identification tests prior to the scan. Participants were paid $20 per hour and provided informed consent to a protocol approved by the Princeton University Institutional Review Board.

### Stimuli

The primary stimulus set included 24 fractal-like images that were masked to be either square or diamond in shape. An additional 144 unique fractal and phase-scrambled images were included in a localizer to identify V1/V2 voxels reliably responsive to the experimental stimuli. All fractal images were created using ArtMatic Pro (www.artmatic.com). Both square- and diamond-masked stimuli subtended ∼4° of visual angle in diameter on the training/testing laptop computer, and 4.5° in the scanner. We counterbalanced the assignment of images to 3-day delay and no delay conditions and to sequences containing either predictive or nonpredictive actions, and randomly assigned images to serve as cues or outcomes. The Psychophysics Toolbox^41^ for MATLAB (MathWorks) was used for stimulus presentation and response collection.

### First training session (3-day delay)

The first training session was proctored three days before the fMRI scan on a laptop computer in a behavioral testing room. As in previous studies involving the same action-based training paradigm, the training session began with an exploratory training phase, followed by a verbal outcome-identification test, and finally a directed training phase^3,42^. The exploratory training phase included 320 trials in which a cue stimulus appeared on the computer screen for 1,000 ms and then a double-headed arrow appeared below the cue. Participants were allowed an unlimited amount of time for each trial to make either a left button-press or a right button-press, in order to replace the cue with an outcome stimulus that appeared for 1,000 ms. A meter at the bottom of the screen tracked the proportion of left and right button presses throughout the exploratory training phase, and participants were instructed to keep the meter pointer within a specified central zone, in order to roughly equate the frequency of actions and outcomes.

The directed training phase included 160 trials in which the onset of the cue was followed by a single-headed arrow that instructed participants to make a left or right button-press for that trial. Directed training was included in order to equate the stimulus frequencies and transitional probabilities of the two outcomes associated with each cue throughout training. For example, if participants responded left more than right during the exploratory training, they were more likely to be instructed to respond right in the directed training.

### Second training session (no delay)

The second training session was also proctored on a laptop computer in a behavioral testing room, but immediately before the fMRI scan and with new cue and outcome stimuli. To minimize interference between stimulus sets from different sessions, we masked one set with squares and the other with diamonds, with the order counterbalanced across participants. The structure of the second training session was identical to the first, with an exploratory training phase, then a verbal outcome-identification test, and finally a directed training phase.

### Predictive actions

For half of the sequences within each training session, actions were highly predictive of the outcome. For instance, given the predictive cue A, outcome B appeared with 95% probability when the left button was pressed, and outcome C appeared with just 5% probability. Similarly, when the right button was pressed, outcome C appeared with 95% probability and outcome B appeared with just 5% probability. Within each training session, participants were exposed to two different cue stimuli for which actions were highly predictive of outcomes.

### Nonpredictive actions

Randomly intermixed with the predictive action trials, the remaining half of the sequences within each training session contained nonpredictive actions: the two outcomes for each cue appeared with equal probability, irrespective of which button was pressed. That is, given the nonpredictive cue D, outcome E or outcome F appeared with 50% probability when either the left or right button was pressed. Within each training session, participants were exposed to two different cue stimuli for which actions were nonpredictive of outcomes.

### Scan task

The task in the fMRI scanner resembled the training sessions. Participants were instructed to continue to keep track of probabilistic relationships between button presses and fractal pairs while in the scanner and they knew to expect a final set of behavioral tests after the scan. A total of 320 sequence trials were organized into eight 6-minute runs. Each run contained sequences from either the first training session or the second training session, alternating between runs. Within each run, four blocks of predictive actions alternated with four blocks of nonpredictive actions. Pairs of runs for each participant contained the same stimuli and block order, with the trial order randomized. Each block included five trials and lasted 22.5 s, followed by 18 s of fixation. To match the outcome probabilities during the scan with the trained probabilities, participants were instructed to balance their left and right responses, and one block of predictive actions in each run contained a trial with an incorrectly predicted outcome (modeled separately and excluded from analysis).

As during exploratory training, each trial in the scanner involved three parts: a cue stimulus for 1000 ms, an action prompt consisting of a double-headed arrow below the cue that remained on screen until a button press or until the 1500 ms response window elapsed, and an outcome stimulus for 1000 ms. Participants used a separate response box for each hand to press the left and right buttons. If participants did not press a button within the response window, the cue stimulus and action prompt were replaced with a fixation cross that remained on screen until the next trial.

### Verbal tests

Each participant performed six verbal tests over the course of the study: one test during each of the two training sessions (between the exploratory and directed training phases), one pre-scan test for each stimulus set directly before the fMRI scan, and one post-scan test for each stimulus set directly after the scan. On each test trial, a cue stimulus appeared at fixation. Below the cue, a single-headed arrow pointed either left or right, and participants were instructed to press the corresponding button. The cue and arrow disappeared, replaced by the two possible outcomes for that cue, presented above and below where the cue had been. For predictive actions, one outcome correctly completed the cue-action-outcome sequence, while the other outcome completed the cue-action-outcome sequence for the other action. Each verbal test included 16 trials (two trials for each cue-action-outcome sequence) with predictive and nonpredictive actions intermixed in a random order. Participants spoke aloud either “top” or “bottom” to indicate which outcome was expected. Verbal responses were used to avoid the button presses used for the trained actions. If a participant was less than 100% accurate for predictive actions in a verbal test, they were allowed to repeat the test one time without receiving feedback about which trials were answered incorrectly. Among the 24 scanned participants, seven participants repeated the pre-scan test for associations from the first training session, and one participant repeated the post-scan test for associations from the second training session. Two additional participants completed the training sessions but did not participate in the fMRI scan because their accuracy was less than 100% upon repeating a pre-scan test.

### Choice RT

Throughout training and in the scanner, we measured choice RT as the time it took for participants to press the left or right button in response to a cue. Although there was an unlimited amount of time for participants to respond during the exploratory and directed training phases, the response window was limited to 1500 ms after the action prompt in the scanner. To make the interpretation of choice RT comparable across the different parts of the experiment, we excluded trials from training in which choice RT exceeded a cutoff of 1500 ms.

### MRI acquisition

Structural and functional MRI data were collected on a 3T Siemens Skyra scanner with a 20-channel head coil. Structural data included a T1-weighted magnetization prepared rapid acquisition gradient-echo (MPRAGE) sequence (1 mm isotropic) for registration and segmentation of EVC, and two T2-weighted turbo spin-echo (TSE) sequences (0.44 × 0.44 × 1.5 mm) for hippocampal segmentation. Functional data consisted of T2*-weighted multi-band echo-planar imaging sequences with 42 oblique slices (16° transverse to coronal) acquired in an interleaved order (1,500 ms repetition time (TR), 40 ms echo time, 1.5 mm isotropic voxels, 128 × 128 matrix, 192 mm field of view, 71° flip angle, acceleration factor 3, shift 2). These slices produced only a partial volume for each participant, parallel to the hippocampus and covering the temporal and occipital lobes. Collecting a partial volume instead of the full brain allowed us to maximize spatial and temporal resolution over our *a priori* ROIs. Data acquisition in each functional run began with 6 s of rest to approach steady-state magnetization. A B0 field map was collected at the end of the experiment.

### fMRI preprocessing

Preprocessing was conducted using the Oxford Centre for Functional MRI of the Brain (FMRIB) Software Library 5.0 (FSL)^43^. Functional runs were corrected for slice-acquisition time and head motion, high-pass temporally filtered using a 128 s period cutoff, spatially smoothed using a 3 mm FWHM Gaussian kernel, and registered to each participant’s high-resolution MPRAGE image using FLIRT boundary-based registration with B0-fieldmap correction^44^.

### Hippocampal segmentation

We anatomically defined hippocampal subfields on high-resolution T_2_-weighted images for each participant, using the automatic segmentation of hippocampal subfields (ASHS) machine learning toolbox^45^ and a database of manual medial temporal lobe segmentations from a separate set of 51 participants^46,47^. These manual segmentations were in turn based on anatomical landmarks from prior studies^48,49^. The hippocampal ROI was formed by combining CA2/3, dentate gyrus, and CA1. This was planned *a priori* because these subfields were linked to pattern completion during action-based prediction in our previous study^3^.

### Early visual cortex

The EVC ROI for each participant was anatomically constrained to V1 and V2, and functionally constrained to voxels reliably responsive to the experimental stimuli, as determined by an independent functional localizer. V1 and V2 were defined in each participant’s T1-weighted anatomical scan using anatomical masks^50,51^ generated with FreeSurfer^52^. During the functional localizer scan, participants detected one-back repetitions of fractals and scrambled images that had been masked to the same shapes (square, diamond) and sizes as the experimental stimuli. Fractal and scrambled stimuli were arranged into 16 blocks, each 15 s in duration with 9 s fixation between blocks. Within each block, 10 stimuli were each presented for 1,000 ms followed by 500 ms fixation between trials. In total, the localizer run was ∼7 min in duration, and included 72 unique fractal images, along with 72 phase-scrambled versions of those images. Within the anatomical boundaries of V1 and V2, we selected voxels that were reliably responsive (*p* <.05) to both the square- and diamond-masked stimuli in the localizer.

### Stimulus-evoked activity

A separate GLM containing FIR basis functions was applied to each run of the preprocessed data using FMRIB’s Improved Linear Model (FILM)^43^ with local autocorrelation correction. Each block and subsequent fixation period were modeled by 27 delta functions, one for each TR. Parameter estimates were averaged over TRs 5-19 of each block to capture the peak stimulus-evoked response. This included the stimulus duration shifted forward by 6 s in order to account for the hemodynamic lag. These stimulus-evoked responses were then averaged across voxels in each ROI and converted to percent signal change before combining across runs for each condition.

### ROI background connectivity

Task-specific background connectivity between the hippocampus and EVC was measured through a previously described approach^12,14^. White matter and ventricle activity, along with six motion parameters, were regressed out of the preprocessed BOLD signal timecourses in a GLM for each run that was fit using FILM. Then, we estimated stimulus-evoked BOLD responses with the same FIR procedure above. Critically, the FIR basis functions captured the average timing and shape of the hemodynamic response in each voxel in a data-driven way. Within each run, we *z*-scored the residual (“background”) timeseries, extracted a 34.5 s time window of data for each block (from the start to 12 s after the last trial), and concatenated the data across blocks for each condition. Background connectivity was then measured as temporal Pearson correlations between concatenated background timeseries from each region and averaged across runs for each participant.

### Across-run control correlations

Control analyses correlated background signal across pairs of runs containing the same stimuli and block order. Insofar as we successfully removed stimulus-evoked responses, across-run correlations should be eliminated, even when the same timeseries produce reliable correlations and correlation differences within run. The across-run correlations also quantify the contribution (if any) of stimulus-evoked responses to the residual background connectivity, which can be used as a baseline for within-run measures.

### Voxelwise background connectivity

We performed exploratory analyses using the residual timeseries from bilateral hippocampal and EVC ROIs to calculate background connectivity with all voxels in the partial volume. The reliability of these maps was assessed across participants by registering the correlation maps for each seed ROI and condition to the MNI152 template space, which had been resampled with interpolation to match the resolution of the functional data (1.5 mm isotropic). Nonparametric randomization tests were performed for each voxel’s connectivity using FSL Randomise^53^, and corrected for multiple comparisons with threshold-free cluster enhancement (TFCE), resulting in a family-wise error rate of *p* <.05.

### Time-lagged background connectivity

To examine the temporal dynamics of hippocampal-EVC background connectivity, we measured the temporal cross-correlation of background activity. Specifically, we shifted the time windows for each block either forward or backward to assess temporal precedence. To test for evidence that the hippocampus leads EVC, we computed the within-block Pearson correlations between background signal in the hippocampus from 0–33 s and EVC from 1.5–34.5 s. Likewise, to test for evidence that EVC leads the hippocampus, we computed the within-block correlations between the hippocampal background activity from 1.5–34.5 s and EVC background activity from 0–33 s. To avoid concerns about the relative timing of the BOLD response between region, we are not interested in the overall magnitude of cross-correlations, but rather in modulation of these cross-correlations by experimental condition.

### Multivariate pattern similarity

Pattern similarity in the hippocampus and EVC was measured as Pearson correlations across voxels within each ROI. Multivoxel patterns for each cue-outcome transition were based on parameter estimates of BOLD response amplitude in an event-related GLM for each run. Each GLM was fit using FILM with local autocorrelation correction and six motion parameters as nuisance covariates. A regressor for each cue-outcome transition was constructed by convolving trial onsets and durations with a double-gamma hemodynamic response function, and a separate regressor was included in each GLM to account for predictive actions with counter-predicted outcomes and trials for which the participant failed to press a button before the response deadline. Parameter estimates for each cue-outcome transition were averaged across runs before calculating pattern similarity. Within-cue pattern similarity for each condition was measured as the correlation between cue-outcome transitions that shared the same cue stimulus but contained different outcome stimuli. Across-cue pattern similarity for each condition was measured as the correlation between cue-outcome transitions containing completely distinct stimuli (different cue and different outcome).

### Statistics

All correlation coefficients calculated for background connectivity and pattern similarity were Fisher *z*-transformed prior to statistical analysis. In ROI analyses, pattern similarity and background connectivity were calculated separately for each hippocampal hemisphere and then averaged across hemispheres to reduce multiple comparisons. Repeated-measures ANOVAs and paired-sample *t*-tests were used to compare background connectivity and pattern similarity for predictive and nonpredictive actions. Tests were evaluated against a two-tailed *p*<0.05 level of significance.

### Data availability

All data and code will be made available freely upon request.

## Acknowledgements

This work was supported by NIH grants F32 EY023162, R01 EY021755, and R01 MH069456.

## Author contributions

N.C.H., E.W.A., and N.B.T.-B. reviewed the analyses, discussed the results, and wrote the paper. N.C.H. and N.B.T.-B. designed the experiment. N.C.H. and E.W.A. collected the data and performed the analyses.

